# Mining signaling flow to interpret mechanisms of synergy of drug combinations using deep graph neural networks

**DOI:** 10.1101/2021.03.25.437003

**Authors:** Heming Zhang, Yixin Chen, Philip Payne, Fuhai Li

## Abstract

Complex signaling pathways/networks are believed to be responsible for drug resistance in cancer therapy. Drug combinations inhibiting multiple signaling targets within cancer-related signaling networks have the potential to reduce drug resistance. Deep learning models have been reported to predict drug combinations. However, these models are hard to be interpreted in terms of mechanism of synergy (MoS), and thus cannot well support the human-AI based clinical decision making. Herein, we proposed a novel computational model, *DeepSignalingFlow*, which seeks to address the preceding two challenges. Specifically, a graph convolutional network (GCN) was developed based on a core cancer signaling network consisting of 1584 genes, with gene expression and copy number data derived from 46 core cancer signaling pathways. The novel up-stream signaling-flow (from up-stream signaling to drug targets), and the down-stream signaling-flow (from drug targets to down-stream signaling), were designed using trainable weights of network edges. The numerical features (accumulated information due to the signaling-flows of the signaling network) of drug nodes that link to drug targets were then used to predict the synergy scores of such drug combinations. The model was evaluated using the NCI ALMANAC drug combination screening data. The evaluation results showed that the proposed *DeepSignalingFlow* model can not only predict drug combination synergy score, but also interpret potentially interpretable MoS of drug combinations.

## 1. Introduction

Complex signaling pathways and ensuing networks are believed to be responsible for drug resistance in cancer therapy. Thus, drug combinations inhibiting multiple signaling targets in cancer-related signaling networks have the potential to be effective in terms of reduce drug resistance. However, it is challenging to discover effective and synergistic drug combinations in clinical and laboratories settings because of the unclear and complex mechanism of synergy (MoS) for such drug combinations. Recent analyses of reported drug combinations indicate that there are complex, diverse and heterogeneous MoS for different drug combinations in different diseases^1^. Further, in recent studies, the RAS/ERK signaling pathway, which has been show to inhibit the efficacy of certain anti-cancer drugs was found to be synergistic with autophagy inhibitors in RAS-driven cancers^2,3^. This is notable, given that autophagy signaling was activated after treatment with RAS inhibitors^2,3^. Despite this evolving body of research concerning the targeting of synergistic cancer therapeutics, there remain few effective drug combinations that are used broadly in the clinical environment. Therefore, novel and effective drug combinations are needed in order to expand the scope and impact of precision cancer therapy and to ultimately improve cancer treatment outcomes.

In response to the aforementioned challenges, and to help facilitate the discovery of synergistic drug combinations for use in a variety of cancer, we sought to understand the associations between genetic biomarkers and drug response, as well as mechanism of drug synergy, using computational methods. The National Cancer Institute (NCI) has previously generated a comprehensive screening data set of ~5,232 combinations from ~100 drugs in 60 human tumor cell-lines (NCI-60), which is publicly available as the NCI-ALMANAC Drug Combination Data Set^4^. This dataset is valuable for developing computational models for drug combination prediction. There have been a variety of computational methods described in the scientific literature concerning the application of machine learning techniques to such data. For example, matchmaker^5^ was developed to analyze chemical and gene expression profiles using a deep learning framework, which is similar as the DeepSynergy^6^ and AuDNNsynergy^7^ models. Similarly, a simplified deep learning model, DeepSignalingSynergy, has been proposed to investigate the discriminative importance of individual signaling pathways for drug combination prediction^8,9^. In addition, the use of multi-task learning with transfer learning model has been proposed to prediction drug combinations^10^. Further, random forest (RF) and XGBoost models have been used, based on the chemical and genomics features of source data, to predict drug combinations^11^. These and many other computational models have also been proposed for drug combination prediction (as opposed to predicting the efficacy of single drugs). For example, the connectivity map (CMAP)^12,13^ and associated network analysis based drug combination prediction models have been used in this context^14,15^. In an analogous manner, message propagation-based models have been developed based on the confluence of drug targeting and genomics data^16,17^. A common thread in the preceding work has been the use of semi-supervised learning models, applied to multiple pharmacogenomics datasets, for drug combination prediction^18^.

However, despite the promise of the existing body of work in this domain, there remain three limitations of the existing deep learning models. First, a large number of genomic and chemical features have been used in such predictive models. Given the limited amount of drug screening data currently available, the training of such models with a large number of features is intrinsically limited. Second, the signaling interactions in gene regulatory signaling networks are often not incorporated in the deep learning models, despite their criticality in terms of treatment efficacy. Third, complex mechanism of synergy (MoS) are usually not modeled in these existing constructs. As such, these models are hard to interpret and explain in order to identify effective and synergistic drug combinations for subsequent verification and validation.

In the specific context of cancer studies, signaling pathways playing important roles in terms of tumor development and drug response, are of critical importance when seeking to identify single or synergistic therapeutic targets. For example, in one set of studies, 10 signaling pathways were systematically analyzed cross 9,125 tumor samples in 33 cancer types (64 subtypes)^19^. The results of this study indicated that 89% of tumor samples had at least one driver alteration in one of these pathways, and 57% and 30% had one and multiple potentially druggable targets, respectively. These results indicate the importance of identifying or repurposing effective drugs and drug combinations that target these signaling pathways. Therefore, in our study, we proposed a novel computational graph neural network (GNN) model, *DeepSignalingFlow*, to investigate the aforementioned issues as they relate to the used of deep learning models for drug combination therapy predication, by incorporating core cancer signaling pathways into such analyses.

## 2. Methodology

### 2.1 NCI ALMANAC drug combination screening dataset

As was introduced previously, the NCI Almanac dataset includes the combo-scores^1^ of permutations of 104 FDA approved drugs relative to the tumor growth of NCI60 human tumor cell lines. The average combo-score of two drugs with different doses on a given tumor cell line was used as the basis for assessing the synergy score of two drugs on the given tumor cell line using a 4-element tuple: <*D*_*A*_, *D*_*B*_, *C*_*C*_, *S*_*ABC*_>.

### 2.2 Gene expression and copy number data of NCI-60 Cancer Cell Lines

Multi-omics data (e.g., RNA-seq [gene expression], copy number variation, metabolomics, miRNA, and RPPA_ for 1000 human tumor cell lines were available in the Cancer Cell Line Encyclopedia (CCLE) database^20^, which are of value when seeking to identify associations between genetic biomarkers and drug response. In addition, copy number data was downloaded from GDSC database^21^. By identifying overlapping cell lines from these two databases, we identified 43 of NCI-60 cancer cell lines that were also present in CCLE (see **Table 1**).

**Table 1:** 43 NCI-60 human tumor cell lines included in CCLE database.

|  |  |  |  |  |
| --- | --- | --- | --- | --- |
| A498_KIDNEY | A549_LUNG | ACHN_KIDNEY | BT549_BREAST | CAKI1_KIDNEY |
| HOP62_LUNG | HOP92_LUNG | HS578T_BREAST | HT29_LARGE_INTESTINE | IGROV1_OVARY |
| K562_HAEMATOPOIETIC_AND_LYMPHOID_TISSUE | KM12_LARGE_INTESTINE | LOXIMVI_SKIN | MCF7_BREAST | MDAMB231_BREAST |
| MDAMB468_BREAST | NCIH226_LUNG | NCIH23_LUNG | NCIH322_LUNG | NCIH460_LUNG |
| NCIH522_LUNG | NIHOVCAR3_OVARY | OVCAR4_OVARY | OVCAR8_OVARY | PC3_PROSTATE |
| RPMI8226_HAEMATOPOIETIC_AND_LYMPHOID_TISSUE | SF268_CENTRAL_NERVOUS_SYSTEM | SF295_CENTRAL_NERVOUS_SYSTEM | SF539_CENTRAL_NERVOUS_SYSTEM | SKMEL28_SKIN |
| SKMEL5_SKIN | SKOV3_OVARY | SW620_LARGE_INTESTINE | T47D_BREAST | U251MG_CENTRAL_NERVOUS_SYSTEM |
| UACC257_SKIN | UACC62_SKIN | UO31_KIDNEY |  |  |

### 2.3 KEGG signaling pathways and cellular process

46 signaling pathways (45 “signaling pathways” + cell cycle)^8,9^ were collected from the KEGG (Kyoto Encyclopedia of Genes and Genomes)^22^ database, specifically: MAPK, FoxO, TGF-beta, T cell receptor, Adipocytokine, ErbB, Sphingolipid, VEGF, B cell receptor, Oxytocin, Ras, Phospholipase D, Apelin, Fc epsilon RI, Glucagon, Rap1, p53, Hippo, TNF, Relaxin, Calcium, mTOR, Toll-like receptor, Neurotrophin, AGE-RAGE, cGMP-PKG, PI3K-Akt, NOD-like receptor, Insulin, Cell cycle, cAMP, AMPK, RIG-I-like receptor, GnRH, Chemokine, Wnt, C-type lectin receptor, Estrogen, NF-kappa B, Notch, JAK-STAT, Prolactin, HIF-1, Hedgehog, IL-17, Thyroid hormone signaling pathways. In these 46 signaling pathways^8,9^, 1584 genes were identified for which gene expression (TPM) and copy number data were available in the CCLE database.

### 2.3 Drug-Target interactions from DrugBank database

Drug-target information were extracted from the DrugBank^23^ database (version 5.1.5, released 2020-01-03). In total, 15,263 drug-target interactions were obtained for 5435 drugs/investigational agents and 2775 targets. Among these drug, 67 drugs were included in NCI ALMANAC with associated targets. Further, 21 drugs with known targets corresponding to the 1584 signaling pathways previously identified were selected for use in our model, specifically: Celecoxib, Gefitinib, Quinacrine hydrochloride, Tretinoin, Cladribine, Imatinib mesylate, Romidepsin, Vinblastine sulfate (hydrate), Dasatinib, Lenalidomide, Sirolimus, Vorinostat, Docetaxel, Mitotane, Sorafenib tosylate, Thalidomide, Everolimus, Nilotinib, Tamoxifen citrate, Paclitaxel, Fulvestrant.

**Figure 1:**
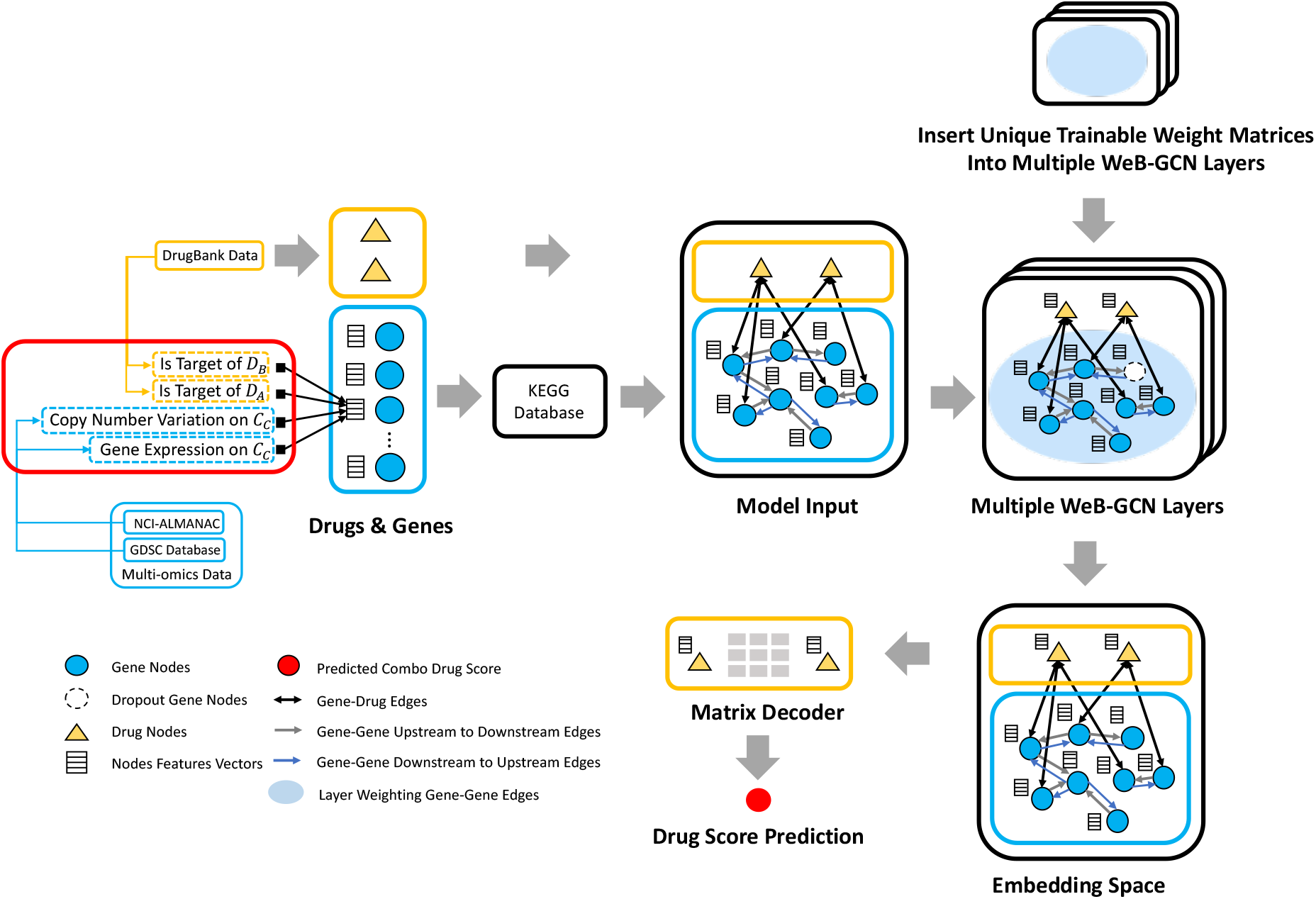
Architecture of DeepSignalingFlow.

### 2.4 Architecture design of DeepSignalingFlow

**Fig.1** shows the schematic architecture of the proposed DeepSignalingFlow model. The model input parameters are: *X* ∈ ℝ^*n*×*n*^, *A* ∈ ℝ^*n*×*n*^, *D*_*in*_ ∈ ℝ^*n*×*n*^, *D*_*out*_ ∈ ℝ^*n*×*n*^, where *X* denotes the nodes features matrix with *n* nodes of *d* features, *A* is the adjacent matrix that links gene-gene interactions on the signaling pathways and gene-drug interactions, and the elements of adjacent matrix *A* such as *a*_*ij*_ indicates an edge from *i* to *j*. *D*_*in*_ is an in-degree diagonal matrix for nodes in directed graph, and *D*_*out*_ is an out-degree diagonal matrix for nodes in in directed graph.

In the graph convolution stages of our architecture, we added the bidirectional nodes-flow with both of ‘upstream-to-downstream’ (from up-stream signaling to drug targets) and ‘downstream-to-upstream’ (from drug targets to down-stream signaling). Therefore, we call this model a Weight Bi-directional Graph Convolution Network (WeB-GCN). The weight matrices are denoted as: 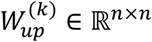,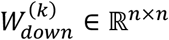(*k* = 1,2,…, *K*) respectively. And then the weight adjacency matrices will be denoted as:

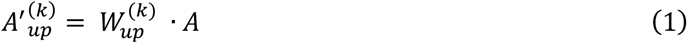

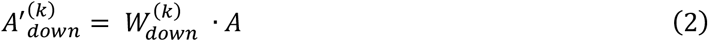

where for each 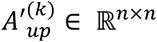,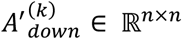(*k* = 1,2,…, *K*) parameters for drug-gene relations values are the same with those in original adjacent matrix. Complete model parameters are specified in **Table 2**.

**Table 2:** Model parameters and notations.

| Input Matrices | When $k = 2, 3, \dots, K$ | When $k = 1$ |
| --- | --- | --- |
| Nodes Features Matrix | $H^{(k-1)} \in \mathbb{R}^{n \times 4d^{(k-1)}}$ | $H^{(0)} \in \mathbb{R}^{n \times d^{(0)}}$ |
| New Nodes Adjacent Matrices | $A'_{up}^{(k)} \in \mathbb{R}^{n \times n}, A'_{down}^{(k)} \in \mathbb{R}^{n \times n}$ | same |
| Nodes In-degree Diagonal Matrix | $D_{in} \in \mathbb{R}^{n \times n}$ | same |
| Nodes Out-degree Diagonal Matrix | $D_{out} \in \mathbb{R}^{n \times n}$ | same |
| Trainable Feature Transformation Matrix | $M_{up}^{(k)} \in \mathbb{R}^{4d^{(k-1)} \times d^{(k)}}$<br>$M_{down}^{(k)} \in \mathbb{R}^{4d^{(k-1)} \times d^{(k)}}$ | $M_{up}^{(1)} \in \mathbb{R}^{d^{(0)} \times d^{(1)}}$<br>$M_{down}^{(1)} \in \mathbb{R}^{d^{(0)} \times d^{(1)}}$ |
| Trainable Biased Feature Transformation Matrix | $B^{(k)} \in \mathbb{R}^{4d^{(k-1)} \times d^{(k)}}$ | $B^{(1)} \in \mathbb{R}^{d^{(0)} \times d^{(1)}}$ |

**Table 2:** 5-fold Cross Validation on 2 Sets of Hyper-parameters.

| Number of Dataset | Input Dims, Output Dims |  |  | Decoder dims | Dropout | Training MSE Loss | Training Pearson Correlation | Test MSE Loss | Test Pearson Correlation |
| --- | --- | --- | --- | --- | --- | --- | --- | --- | --- |
|  | 1<sup>st</sup> layer | 2<sup>nd</sup> layer | 3<sup>rd</sup> layer |  |  |  |  |  |  |
| 1<sup>st</sup> dataset | (4, 12) | (12, 12) | (12, 36) | 40 | 0.01 | 34.11 | 74.64% | 53.34 | 68.20% |
| 2<sup>nd</sup> dataset | (4, 12) | (12, 12) | (12, 36) | 40 | 0.01 | 33.48 | 74.40% | 51.91 | 64.21% |
| 3<sup>rd</sup> dataset | (4, 12) | (12, 12) | (12, 36) | 40 | 0.01 | 33.01 | 75.75% | 44.59 | 66.64% |
| 4<sup>th</sup> dataset | (4, 12) | (12, 12) | (12, 36) | 40 | 0.01 | 32.78 | 75.71% | 42.19 | 67.93% |
| 5<sup>th</sup> dataset | (4, 12) | (12, 12) | (12, 36) | 40 | 0.01 | 35.38 | 73.39% | 48.01 | 62.48% |
| Average | (4, 12) | (12, 12) | (12, 36) | 40 | 0.01 | **33.76** | **74.78%** | **48.01** | **65.89%** |

**Figure 2:**
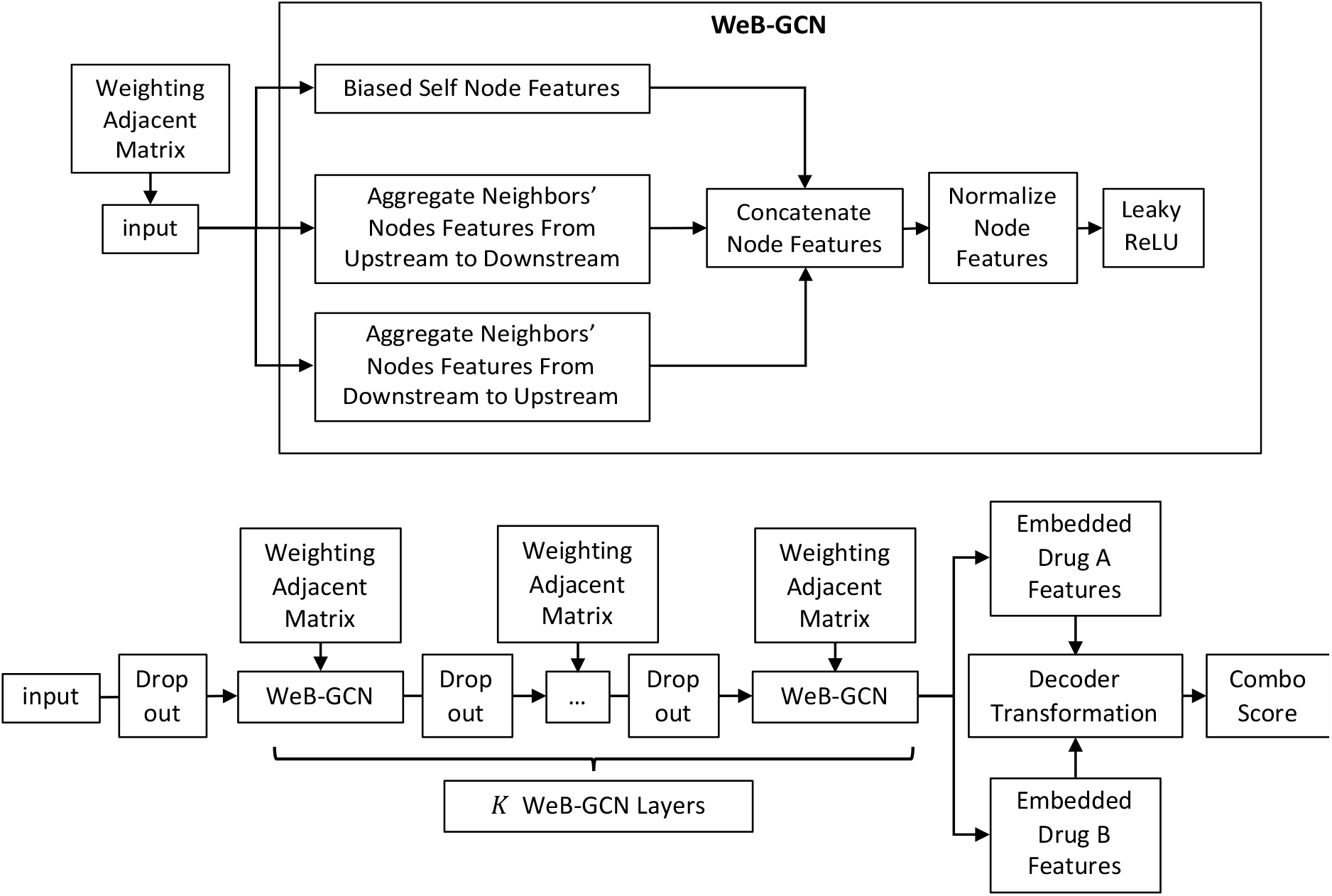
Schematic architecture of the DeepSignalingFlow Model.

**Fig. 2** shows the structural details of our model. First, the mean aggregation of each node neighbors’ features from upstream to downstream is achieved by:

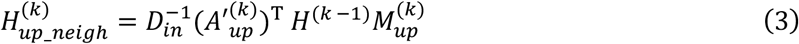

And the mean aggregation of each node neighbors’ features from downstream to upstream is achieved by:

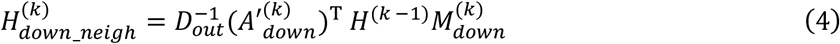

And biased transformation of nodes themselves and their features is achieved by:

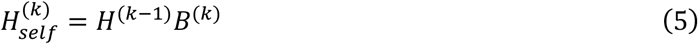

For *k* = 1,2,…, K, the model uses normalization function *N*: 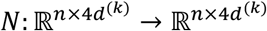and LeakyReLU function with parameter α to map concatenated nodes features 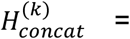 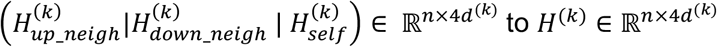 with the equation:

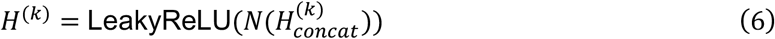

where normalization function *N* enable L2 normalization for demo matrix *V* ∈ ℝ^*p*×*q*^ in row axis with the equation:

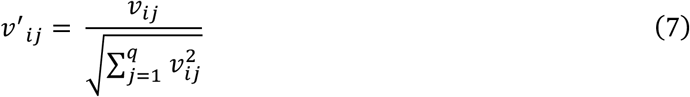

where ν′_*ij*_ is the element of new matrix *V*′ ∈ ℝ^*p*×*q*^. Therefore, the output of *k*-th layer WeB-GCN is *H*^(*k*)^. After obtaining the embedded nodes features with 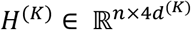, the embedded nodes features of drug_A are obtained: 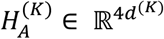, and the drug_B embedded nodes features are obtained as: 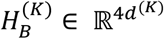, which are just the 1585^th^ and 1586^th^ rows in *H*^(*k*)^. Based on the idea of decagon decoder^24^, the prediction of combo score will be calculated in following equation:

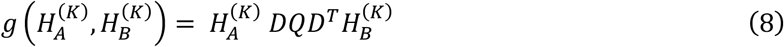

where 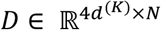 and *D* ∈ ℝ^*N*×*N*^ are trainable decoder matrices.

### Drug combination response interpretation using the directional signaling flow

As aforementioned, we added directional signaling interaction weights to indicate the potential signaling flow from up-stream signaling to drug targets, and from drug targets to down-stream signaling. The trainable directional weights can potentially indicate and interpret the signaling flow on the signaling network to affect the drug combination response. Specifically, the signaling flow derived from the directional weighted matrices are defined as follows. Each such model had k=3 layers, and each layer has upstream weight adjacent parameters and downstream weight adjacent parameters, which are 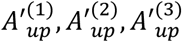and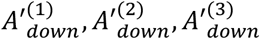. Through aggregating 3 layers upstream and downstream to weight adjacent matrices, we arrive at an aggregated upstream, downstream and bind-stream for *k*th 5-fold cross valuation model with *k*th model’s
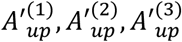and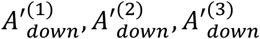 using the formula:

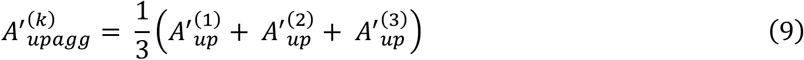

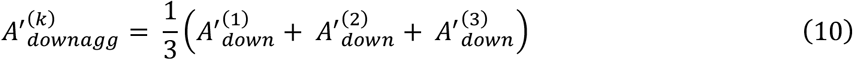

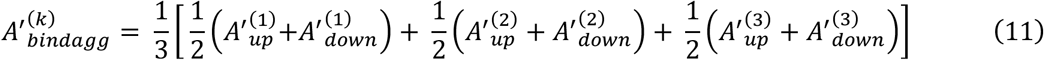

With upstream, downstream and bind-stream weight adjacent matrices parameters, we then aggregated weight adjacent matrices of those 5 models by the following formula:

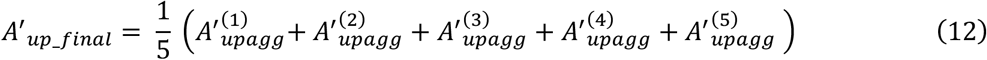

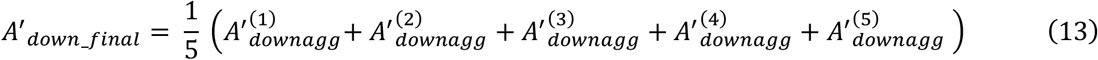

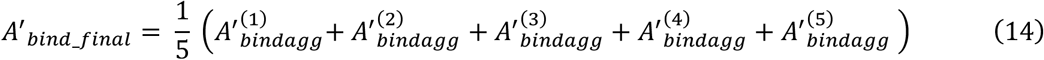

With the final upstream, downstream and bind-stream weight adjacent matrices parameters for 1584 genes of *A*′_*up_final*_, *A*′_*down_final*_, *A*′_*bind_final*_, we then were able to study the signaling flow on the signaling network.

## 3. Results

### 3.1 Experimental Setup

Model input: <drug_A, drug_B, cellline name, combo score> were used to construct each point in an input dataset, which was comprised of 5445 such points; among these data, using a 5-fold cross validation with 5 splits of 1089 nodes, we created a training dataset. For each point, the cell line name contained the RNA sequence of the 1584 genes. Given all possible 2-drug permutations, each point created a graph with 1586 nodes, and each such graph had 3 initial features. For those 1584 genes, their 3 features indicate the RNA sequence number, and whether they have connection spanning drug_A and drug_B. For each 2-drug dyad, their 3 features were initiated with zeros. Furthermore, to indicate connections between nodes, adjacent matrices were also created. Those matrices were formed from file ‘Selected_Kegg_Pathways_edges_1’, which contains gene pairs with sources and destinations. In this way, the graphs generated were directed graphs. For drug-gene edges, the connections are bidirectional, which means that in adjacent matrices, those elements are symmetric. At the same time, in-degree matrices are formed according to said adjacent matrices.

### 3.2 Hyperparameters Setting

Subsequently, a model was developed by using pytorch. Using the Adam optimizer, learning rate started with 0.01 and reduced equally within each batch of first 50 epochs. We empirically set the training epochs as 75, which allows for optimal validation results. *K* = 3 WeB-GCN layers were used, and the initial input feature dimension was: *d*^(0)^ = 3, with the feature dimensions varying at the different layers, and denoted by: (*d*^(1)^,*d*^(2)^,…, *d*^(*K*)^), *k* = 1,2,3. WeB-GCN concatenate biased last layer node features and transformed nodes features, with the final concatenated dimensions of the WeB-GCN layers for the output dims being 3 × *D*^(*K*)^. Output dims of the last layer served as the input dims for this layer, as follows:

1. First layer (input dims, output dims): (*d*^(0)^, 4*d*^(1)^)
2. Second layer (input dims, output dims): (4*d*^(1)^), 4*d*^(2)^)
3. Third layer (input dims, output dims): (4*d*^(2)^), 4*d*^(3)^)

The final embedded drug nodes dims were 4*d*^(3)^ (see **Table 2**). Decoder trainable transformation matrix dims: 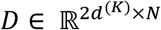 and *Q* ∈ ℝ^*N*×*N*^ were ysed as trainable decoder matrices, with changeable parameters of *N* to adapt model performance. As shown in **Table 2**, the model uses *N* = 40. As for the LeakyReLU function, the parameter *α* = 0.1.

### 3.3. Synergistic score prediction evaluation

To evaluate the model performance in terms of synergy score prediction for drug combinations, we conducted 5-fold validation. As seen, the average prediction (using the Pearson correlation coefficient), was about 66% accuracy using the testing data. These prediction results are comparable with existing deep learning models^6,7^. The results indicated the feasibility of drug combination synergy score prediction using a graph neural network with a small set of core signaling pathways genes, compared with the existing complex deep neural network (DNN) models using a large number of genomics and chemical structure features.

**Figure 3:**
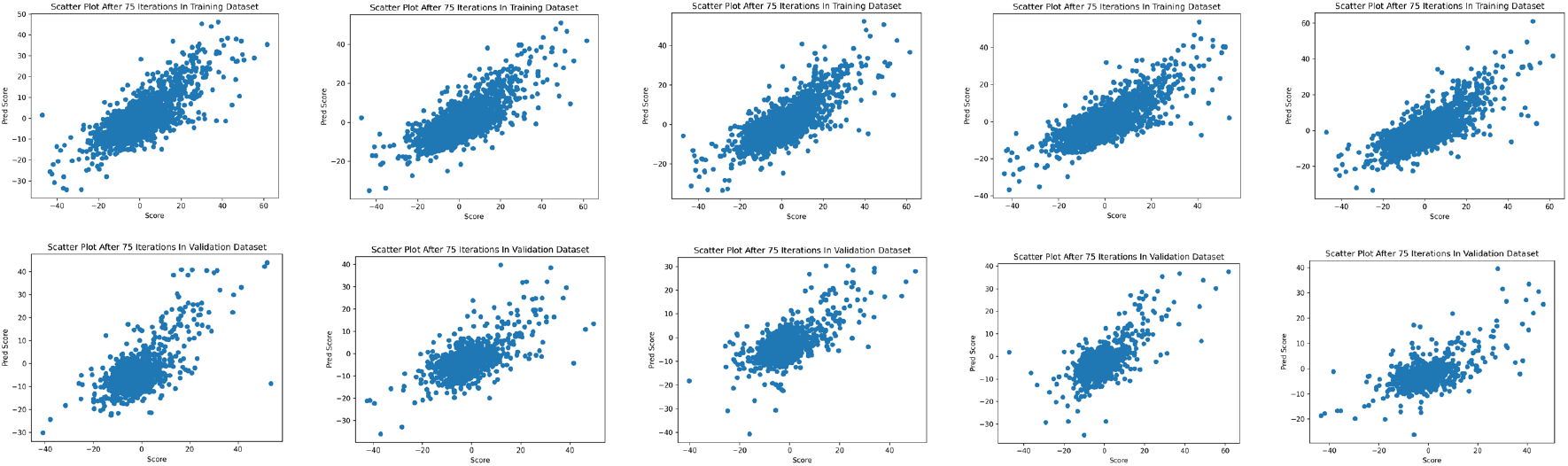
Scatterplots of the predicted and experimental synergy scores on the 5-fold training and validation dataset with dropout rate 0.01

### 3.3 Signaling flow interpreting the mechanism of synergy

As aforementioned, we added directional signaling interaction weights to indicate the potential signaling flow from up-stream signaling to drug targets, and from drug targets to down-stream signaling. In this sub-aim, we identified a set of important sub-network, including important signaling interactions, to show the feasibility of model interpretation. As an example, first, the signaling interactions (edges) were filtered, based on the trained signaling interaction weights, by selecting the signaling interactions with weights more than a given threshed, which indicates the important signaling flow on the signaling network. Then network nodes with degrees larger than 50 were selected to generate signaling flow network to partially explain the mechanism of drug combination synergy (see **Table 3**). Different thresholds of edges and nodes could generate different signaling flow subnetworks. As seen in **Figs. 4,5,6**, the core up-stream, down-stream and integrated (up- and down-stream) signaling flow networks were identified from the large signaling network based on the average trained directional weight matrices on the 5 training datasets. This sub-network module has the potential to uncover the potential mechanism of synergy of the involved drugs.

**Table 3:** Filtering results for upstream

| Network | Filtering Threshold | Nodes Number | Edges Number | Target Genes Number |
| --- | --- | --- | --- | --- |
| Upstream | Nodes Degree > 50 | 105 | 1034 | 20 |
| Downstream | Nodes Degree > 50 | 100 | 1101 | 14 |
| Bindstream | Nodes Degree > 50 | 106 | 1095 | 19 |

**Figure 4:**
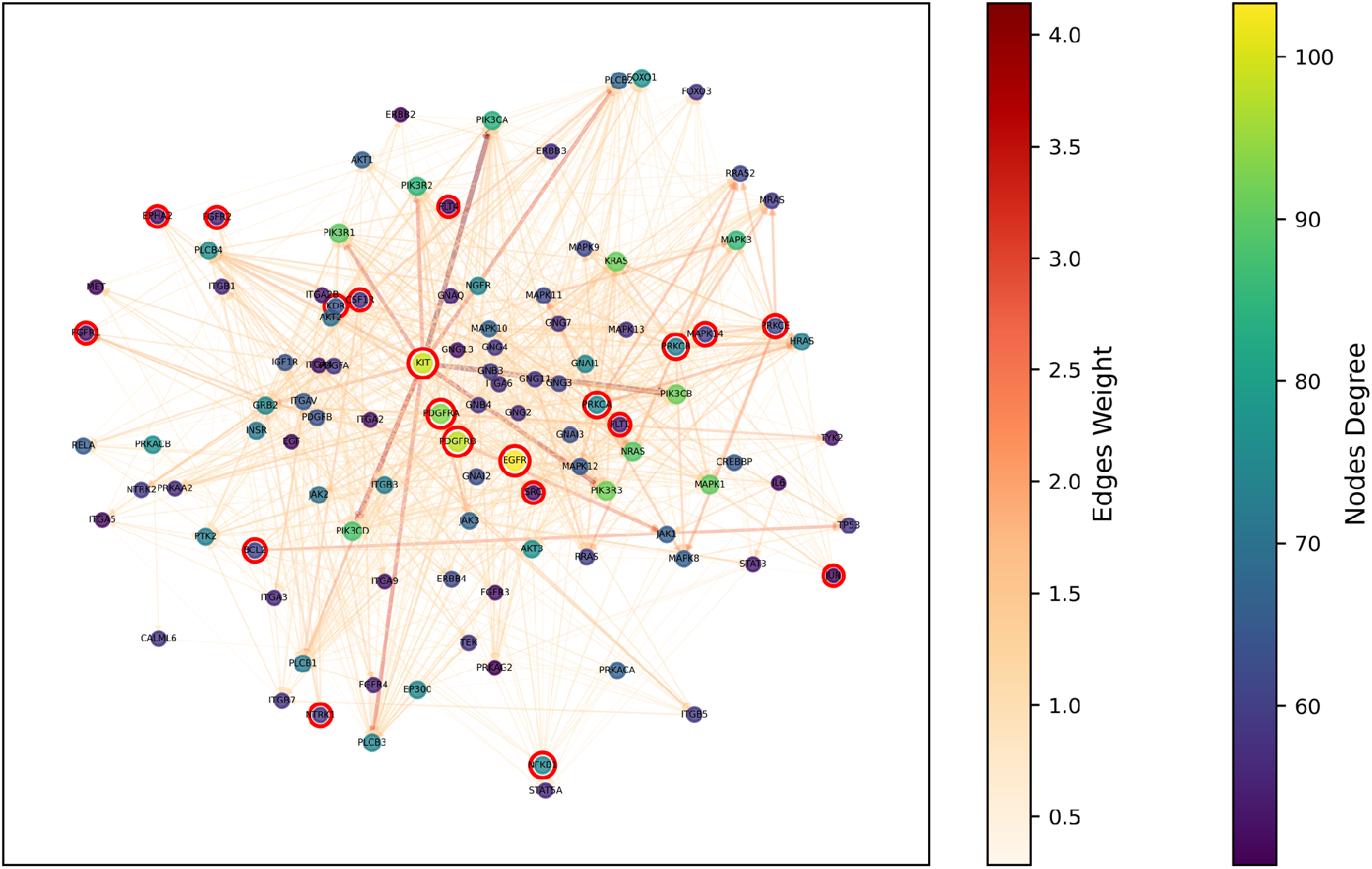
Up-stream core signaling targets and signaling flows among these targets (nodes circled with red are targets of drugs).

**Figure 5:**
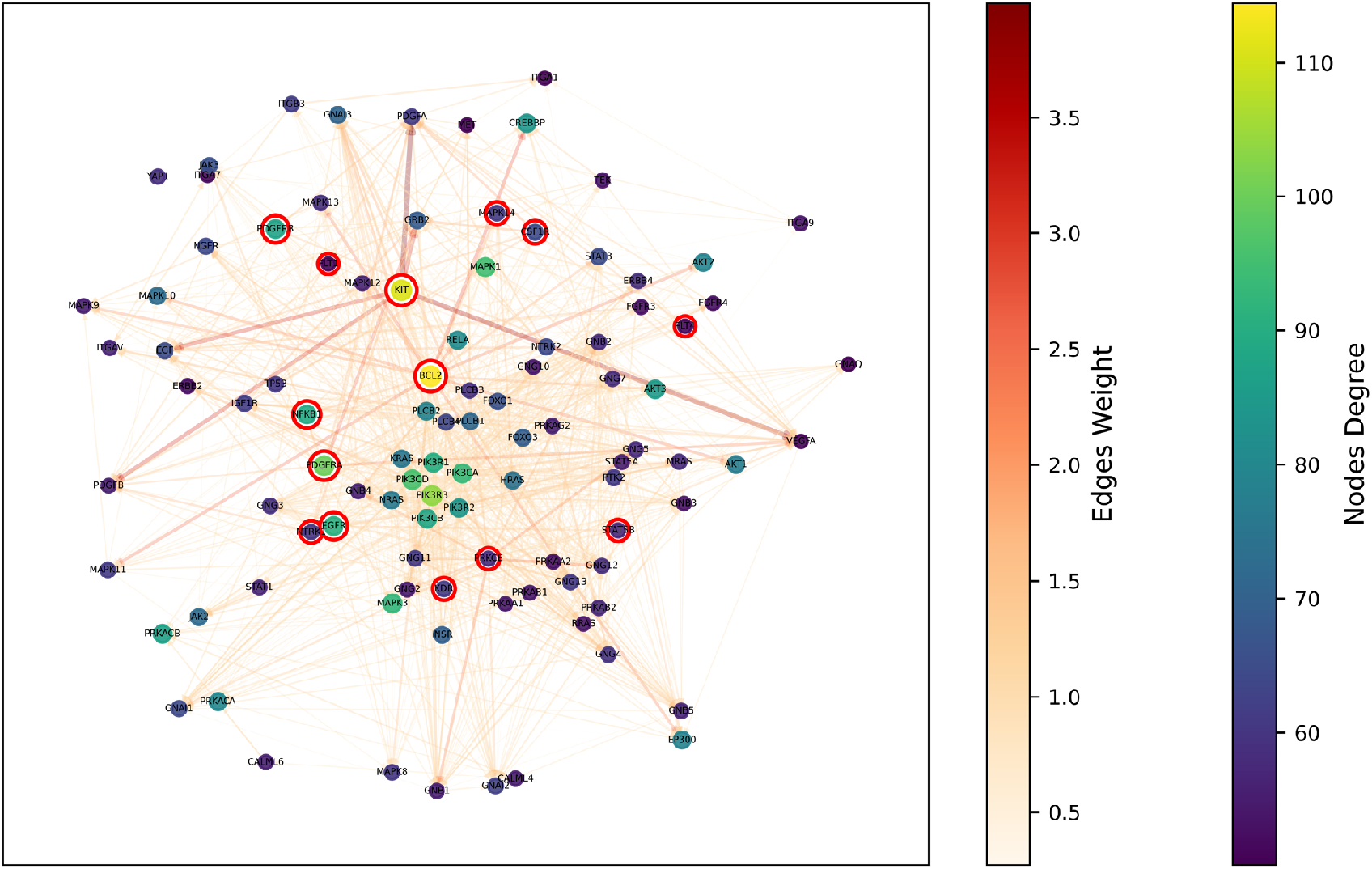
Down-stream core signaling targets and signaling flows among these targets (nodes circled with red are targets of drugs).

**Figure 6:**
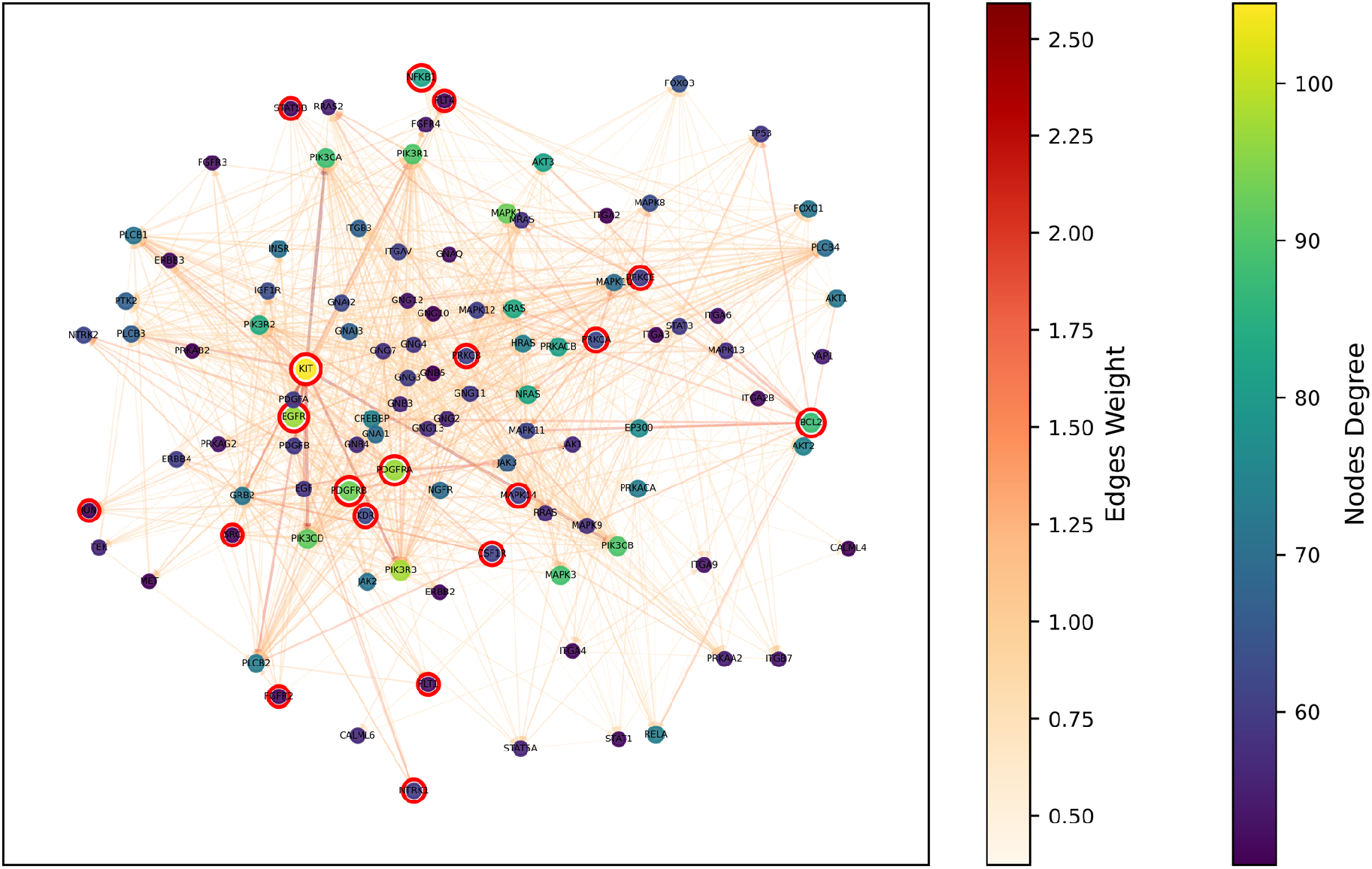
Integrated (up-and down-stream) core signaling targets and signaling flows among these targets (nodes circled with red are targets of drugs).

## 4. Discussion and conclusion

Along with the advancement of next-generation sequencing (NGS) technology, multi-omics data, like gene expression, copy number variation, methylation, genetic mutation, microRNA, are being generated to characterize the dysfunctional molecules and complex signaling pathways in cancer patients. The valuable multi-omics data provides the basis for personalized medicine or precision medicine prediction. Drug combinations are being investigated to reduce the drug resistance influenced by the complex signaling networks in cancer. However, it remains an open problem and challenging because of the mysterious, heterogenous molecular mechanisms of synergy of drug combinations.

Deep learning models have been reported to predict drug combinations by simply concatenating a large number of multi-omics data and chemical structure information using the deep neural network (DNN) model. One limitation of the existing DNN models is the missing of complex and biological meaningful gene regulatory relationships. Thus, it is hard to interpret these models to support the real clinical use, which requires the clinical experts to understand the molecular mechanisms used in the AI prediction.

In this study, we aim to explore the feasibility of building an interpretable AI model using the proposed *DeepSignalingFlow* model. The model is built on a core cancer signaling network consisting of 1584 genes, with gene expression and copy number data derived from 46 core cancer signaling pathways. The novel up-stream signaling-flow (from up-stream signaling to drug targets), and the down-stream signaling-flow (from drug targets to down-stream signaling), were designed using trainable weights of network edges. The selected signaling interaction edges with large weights indicate 1) the important biomarker genes; and 2) the signaling cascades among them, which can potentially interpret the model prediction results. In conclusion, the first-time proposed up-stream and down-stream signaling flow design is novel and can potentially interpret the mechanism of drug combination response.

This is still an exploratory study to prediction and investigate drug combination response using deep neural graph network models. There are some limitations to be further investigated. For example, more signaling pathways and protein-protein interactions should be considered to include more drugs. Moreover, more omics data, like mutation, methylation, miRNA, should be integrated and investigated. In addition, our current analysis is a global analysis or pan-cancer analysis by considering all drugs and all cell lines together. However, the drug combination, and cell line specific signaling flows might be different. Thus, it is interesting to investigate the drug- and cell-specific signaling flows that are informative for the drug combination response prediction. We will investigate these challenges in future work.

